# Adverse outcomes in SARS-CoV-2 infected pregnant mice are gestational age-dependent and resolve with antiviral treatment

**DOI:** 10.1101/2023.03.23.533961

**Authors:** Patrick S. Creisher, Jamie L. Perry, Weizhi Zhong, Jun Lei, Kathleen R Mulka, Hurley Ryan, Ruifeng Zhou, Elgin H. Akin, Anguo Liu, Wayne Mitzner, Irina Burd, Andrew Pekosz, Sabra L. Klein

**Author notes:** To whom correspondence should be addressed: Sabra L. Klein, Andrew Pekosz, & Irina Burd. co-first authors (Patrick S. Creisher, Jamie L. Perry).

## Abstract

SARS-CoV-2 infection during pregnancy is associated with severe COVID-19 and adverse fetal outcomes, but the underlying mechanisms remain poorly understood. Moreover, clinical studies assessing therapeutics against SARS-CoV-2 in pregnancy are limited. To address these gaps, we developed a mouse model of SARS-CoV-2 infection during pregnancy. Outbred CD1 mice were infected at embryonic day (E) 6, E10, or E16 with a mouse adapted SARS-CoV-2 (maSCV2) virus. Outcomes were gestational age-dependent, with greater morbidity, reduced anti-viral immunity, greater viral titers, and more adverse fetal outcomes occurring with infection at E16 (3^rd^ trimester-equivalent) than with infection at either E6 (1^st^ trimester-equivalent) or E10 (2^nd^ trimester-equivalent). To assess the efficacy of ritonavir-boosted nirmatrelvir (recommended for pregnant individuals with COVID-19), we treated E16-infected dams with mouse equivalent doses of nirmatrelvir and ritonavir. Treatment reduced pulmonary viral titers, decreased maternal morbidity, and prevented adverse offspring outcomes. Our results highlight that severe COVID-19 during pregnancy and adverse fetal outcomes are associated with heightened virus replication in maternal lungs. Ritonavir-boosted nirmatrelvir mitigated adverse maternal and fetal outcomes of SARS-CoV-2 infection. These findings prompt the need for further consideration of pregnancy in preclinical and clinical studies of therapeutics against viral infections.

## 1. Introduction

Pregnancy is a risk factor for developing severe COVID-19, with pregnant individuals at increased risk of hospitalization, mechanical ventilation, and mortality compared to non-pregnant patients (1–9). The time of infection during gestation contributes to increased severity, with hospitalization and intensive care unit admission being greater in the third than either the second or first trimester (10, 11). While the specific mechanisms that contribute to the increased risk of severe outcomes during pregnancy are not specified, both immunological and physiological changes are likely involved. The immune system undergoes unique shifts as pregnancy progresses, including increased regulatory T and B lymphocytes as well as reduced cytotoxic and cellular immunity, to protect the developing semi-allogenic fetus (12, 13). The general anti-inflammatory shift during the second and third trimesters also may increase the risk of severe outcomes from viruses, including SARS-CoV-2, by blunting anti-viral immune responses (13). Moreover, physiological changes associated with pregnancy including cardiovascular, respiratory, endocrine, and metabolic alterations may further contribute to disease severity (14). While these pregnancy-associated factors are hypothesized to contribute to severe disease and death following infection with SARS-CoV-2, the exact mechanisms contributing to severe COVID-19 disease during pregnancy in humans remain unknown.

In addition to causing severe outcomes in pregnant patients, SARS-CoV-2 infection during pregnancy also can result in adverse fetal outcomes including preterm birth, stillbirth, small size for gestational age, and reduced birth weight (5, 15–19), as well as increased risks of neurobehavioral deficits and delayed motor skills in infants born to infected mothers (20, 21). Like maternal disease, adverse perinatal and fetal outcomes appear to be influenced by gestational age, with greater risk observed after infection in the third as compared with either the second or first trimester (10, 20, 22). Direct placental infection or vertical transmission of SARS-CoV-2 is exceedingly rare (23–25), and thus is unlikely to be the source of adverse fetal and neonatal outcomes. The exact mechanisms underlying these adverse outcomes remain unknown.

Because of their risk for severe COVID-19 and adverse pregnancy outcomes, pregnant individuals are prioritized for receipt of available emergency use authorized antivirals and vaccines (26–28), despite being excluded from clinical trials of SARS-CoV-2 vaccines and antivirals (29). SARS-CoV-2 mRNA vaccines have been proven to be safe and effective during pregnancy (30–32), and the United States Centers for Disease Control and Prevention recommends vaccination for people who are pregnant, recently pregnant, or trying to become pregnant (30). The safety and efficacy of SARS-CoV-2 antivirals during pregnancy has not been as well studied. In the United States, pregnant people are recommended to receive the antivirals remdesivir (brand name Veklury) and ritonavir-boosted nirmatrelvir (brand name Paxlovid) when indicated (33). While neither antiviral included pregnant people in their clinical trials (34), observational studies of remdesivir indicate its safety and efficacy in pregnant populations (35). Nirmatrelvir is an oral antiviral that inhibits the SARS-CoV-2 M^PRO^ protease and is packaged with ritonavir, a previously established HIV protease inhibitor and pharmacologic booster, which does not have direct antiviral effects on SARS-CoV-2 but instead works to prolong the bioavailability of nirmatrelvir through the inhibition of the hepatic cytochrome P-450 (CYP) 3A4 enzyme (36, 37). Ritonavir-boosted nirmatrelvir treatment during pregnancy appears safe, with no adverse obstetric outcomes reported in small observational studies (38–40). The efficacy of ritonavir-boosted nirmatrelvir in preventing SARS-CoV-2 infection or disease during pregnancy remains an open question, in part because most studies to date were not designed to evaluate efficacy (38–40).

Animal models of microbial infections during pregnancy provide mechanistic insight into adverse maternal and fetal outcomes by enabling deeper analysis of vertical transmission and maternal and fetal immune responses. Animal models have elucidated the pathogenesis of infections such Zika virus, influenza A virus, *Plasmodium falciparum,* and Group B *Streptococcus* infections during pregnancy (26). In the absence of human clinical trials, animal models of infection during pregnancy can be used to characterize the safety and efficacy of therapeutics in this high-risk population. To date, published animal models of SARS-CoV-2 infection during pregnancy have been limited (25, 41), which has hindered investigation into both host and viral factors that may underlie the severe outcomes observed in humans. Animal models have only been used to study the potential reproductive toxicity of nirmatrelvir in rats, rabbits, and zebrafish, with no evidence of embryonic toxicity, fetal abnormalities, maternal toxicity, or other adverse outcomes (42, 43). Whether equivalent dosing of nirmatrelvir administered during pregnancy is equally efficacious against SARS-CoV-2 infection in pregnancy as in nonpregnant animals has not been reported.

In the current study, we developed a mouse model of SARS-CoV-2 infection during pregnancy to investigate maternal and offspring outcomes associated with severe COVID-19 disease during pregnancy and elucidate the contribution of gestational age, pulmonary and placental involvement in adverse outcomes, and control of virus replication. Further, we sought to assess the efficacy of ritonavir-boosted nirmatrelvir in limiting virus replication, preventing maternal disease, and mitigating adverse offspring outcomes. Our results demonstrate that SARS-CoV-2 infection during late gestation causes more severe maternal disease and adverse offspring outcomes than infections earlier during gestion, with maternal disease and adverse offspring outcomes associated with reduced pulmonary anti-viral type 1 interferon (IFN) responses, greater viral replication in the lungs, and loss of placental trophoblasts. Treatment with ritonavir-boosted nirmatrelvir not only reduced pulmonary virus replication, but prevented severe disease and adverse fetal outcomes, highlighting additional benefits of antiviral treatment during pregnancy.

## 2 Materials and Methods

### 2.1 Viruses and cells

A mouse adapted strain of SARS-CoV-2 (maSCV2), originally generated by Dr. Ralph Baric (44) was obtained from the Biodefense and Emerging Infections Research Resources Repository (BEI Resources #NR-55329). The maSCV2 virus was originally generated via infectious clone technology using the sequence of SARS-CoV-2/human/USA/WA-CDC-02982586-001/2020 (WA1 strain) with added mutations in the Spike protein that were predicted to increase binding to murine ACE2(45). This virus was further adapted to mice by sequential passage to generate increased virus replication and disease(44). Working stocks of maSCV2 virus were generated by infecting Vero-E6-TMPRSS2 cells at a multiplicity of infection (MOI) of 0.01 tissue culture infectious dose 50 (TCID_50_) per cell in infection media [Dulbecco’s Modified Eagle Medium (DMEM; Sigma # D5796) supplemented with 2.5% filter-sterilized fetal bovine serum (Gibco# 10-437-028), 100 U/ml penicillin and 100 μg/ml streptomycin (Gibco #15149-122), 1 mM l-glutamine (Gibco #2503081), and 1-mM sodium pyruvate (Gibco #11-360-070)]. Approximately 72 hours post infection, the supernatant fluids were collected, clarified by centrifugation (400g for 10 minutes), and stored in aliquots at −70°C.

### 2.2 Experimental mice

Adult (8-12 weeks of age) timed pregnant and nonpregnant female CD-1 IGS mice were purchased from Charles River Laboratories. Pregnant mice arrived on embryonic day (E) 4, E8, and E14 and were singly housed, and nonpregnant female mice were housed at 5 per cage before and after inoculation. Mice were housed under standard animal biosafety level three (ABSL3) housing conditions with *ad libitum* food and water. Mice were given at least 24 hours to acclimate to the ABSL3 facility prior to infections (46). All monitoring and experimental procedures were performed at the same time each day.

### 2.3 SCV2 infections and monitoring

All animal experiments and procedures took place in an ABSL3 facility at the Johns Hopkins School of Medicine. Experimental pregnant mice were intranasally infected at E6, E10, or E16 with 10^5^ TCID_50_ of maSCV2(44) in 30 μl of DMEM (Sigma #D5796) or mock inoculated with 30 μl of media. Dose-response studies in nonpregnant inbred female mice indicate that maSCV2 requires doses of 10^4^ or 10^5^ TCID_50_ to cause disease in adult mice (44). Prior to intranasal infection, mice were anesthetized via intraperitoneal ketamine/xylazine cocktail (80 mg/kg ketamine, 5 mg/kg xylazine). Following intranasal infections, body mass and clinical signs of disease were monitored once daily in the morning for 14 days or until tissue collection. Clinical scores, determined in the home cage, were administered to mice on a scale of 0-4, with one point given for piloerection, dyspnea, hunched posture, and absence of an escape response on each day (47, 48). Clinical scores over the course of 14 days for each animal were summed to give a cumulative clinical disease score.

### 2.4 Antiviral treatment

Experimental animals were administered vehicle alone [1% (w/v) Soluplus (BASF #50539897), 1% (w/v) Tween 80 (Sigma #59924), 0.5% (w/v) methylcellulose (Sigma #94378) in purified water], high dose nirmatrelvir alone (300mg/kg; MedChem Express #HY-138687), or an animal equivalent dose of nirmatrelvir boosted with ritonavir [1.7 mg nirmatrelvir/dose (MedChem Express #HY-138687), 0.6 mg ritonavir/dose (Sigma #(#155213-67-5)]. Animal equivalent doses were calculated as described (49) by converting the standard human dose of nirmatrelvir and ritonavir (50) to a body-surface-area equivalent for mice (49) using a standardized body surface area for mice of 0.007 mg/m^2^. According to the United Sates Food and Drug Administration, this calculation is recommended for conversion of animal doses to human equivalent doses (51), along with an assumed mass of 30g for all calculations so that pregnant and nonpregnant animals receive the same amount per dose. Mice were administered treatment via oral gavage twice daily for 5 days or until tissue collection, starting 4 hours after infection as described in the original published pre-clinical study of nirmatrelvir (52).

### 2.5 Offspring measurements and behavior

Offspring from mock inoculated dams and maSCV2 infected dams were measured at postnatal day (PND) 0, within 12 hours of birth. Body mass (g), length measured from nose to anus (mm), and head diameter measured from ear to ear (mm) were recorded for each pup, directly, using a caliper, and the average for each independent litter was calculated to avoid confounding litter effects. Pups at PND5 were subjected to developmental neurobehavioral assays of surface righting, cliff aversion, and negative geotaxis as described (53, 54). For each test, 1–2 male and 1–2 female offspring from at least 5 independent litters were used per condition to avoid confounding litter effects. Pups were subjected to 3 attempts at each test, with the time to complete each test recorded on a stopwatch. The upper limit of time was 60 seconds, 30 seconds, and 60 seconds for surface righting, cliff aversion, and negative geotaxis, respectively. The pups’ best trial for each test was used for analysis.

### 2.6 Diffusion capacity of carbon monoxide

To measure lung function of experimental mice, diffusion capacity for carbon monoxide (DF_CO_) was measured. Modifications to a previously published protocol (55) were made for application in ABSL3. Mice were anesthetized with ketamine/xylazine cocktail (80 mg/kg ketamine, 5 mg/kg xylazine). Mice were tracheostomized with an 18-G stub needle. For each mouse, two 3 ml syringes containing 0.8 ml of ∼0.5% neon (Ne, an insoluble inert tracer gas), ∼0.5% carbon monoxide (CO), and balanced air were pre-filled and sealed with a 4-way stop cock. Following tracheostomy, gas was injected into the tracheostomy stub-needle to inflate the lungs for two seconds and held for eight seconds. After eight seconds, the 0.8 ml volume was withdrawn back into the syringe in two seconds and the syringe’s stop cock closed, then the gas in the syringe was diluted to 2 ml with room air and resealed. This was repeated using the second syringe for each mouse. Mice were euthanized via cervical dislocation. The closed syringes were decontaminated in an oven at 75°C for 15 minutes within the ABSL3 to inactivate any virus in the gas sample. DF_CO_ was measured using gas chromatography as previously described (55).

### 2.7 Tissue and serum collection

Experimental dams (infected at E6, E10, or E16) or nonpregnant female mice were euthanized at 3 days post-infection. Mice were anesthetized via isoflurane and exsanguination was preformed via cardiac puncture. At the time of euthanasia, the total number of viable and nonviable fetuses was quantified for each pregnant dam. Fetal viability was determined as the percentage of fetuses within uterine horns that were viable. Fetuses were counted as nonviable if they were smaller or discolored compared to gestational age-matched live fetuses or if a fetus was absent at an implantation site (54, 56, 57). Maternal lungs were collected, separated by lobe, and flash frozen on dry ice for homogenization. The left lung was inflated and fixed in zinc buffered formalin (Thermo Fisher Scientific #NC9351419) for at least 72 hours in preparation for histology. Fetuses and placenta were flash frozen in dry ice or fixed in 4% paraformaldehyde (Thermo Fisher Scientific #J19943.K2) for 72 hours at 4°C for immunohistochemistry. Serum was separated by blood centrifugation at 5,000 rpm for 30 min at 4°C. A subset of uninfected pregnant (E16) and nonpregnant adult mice were euthanized and the median liver lobe was collected and flash frozen in dry ice for Western blot.

### 2.8 Pulmonary histopathology

Fixed lungs were sliced into 3-mm blocks, embedded in paraffin, sectioned to 5 μm, mounted on glass slides, and stained with hematoxylin and eosin (H&E) solution to evaluate lung inflammation. Semiquantitative histopathological scoring was performed by a board-certified veterinary pathologist, blinded to study group assignments and outcomes, to measure both severity of inflammation and the extent of inflammation (58–60). Severity of perivascular and peribronchiolar mononuclear inflammation was scored on a scale of 0- to 4 (0, no inflammation; 1, 1 cell layer; 2, 2-3 cell layers; 3, 4-5 cell layers; 4, > 5 cell layers). Severity of alveolar inflammation was scored on a scale of 0-4 (0, no inflammation; 1-increased inflammatory cells in alveoli, septa clearly distinguished; 2 – inflammatory cells fill alveoli, septa clearly distinguished; 3 – inflammatory cells fill multiple adjacent alveoli, septa difficult to distinguish; 4 – inflammatory cells fill multiple adjacent alveoli with septal necrosis). Extent of inflammation was scored separately for perivascular, peribronchiolar, and alveolar areas on a scale of 0 to 4 (0, no inflammation; 1, 2-25% tissue affected; 2, up to 50% tissue affected; 3, up to 75% tissue affected; 4, >75% of tissue affected). Individual scores were summed to give a cumulative inflammation score.

### 2.9 Infectious virus and viral genome copy number quantification and tissue inactivation

Frozen right cranial lungs, nasal turbinates, placentas, and fetuses were homogenized in lysing matrix D bead tubes (MP Biomedicals #6913100). Homogenization media [500ml DMEM (Sigma # D5796), 5ml penicillin/streptomycin (Gibco #15149-122)] was added to bead tubes containing tissue at a minimum volume of 400μl and maximum volume of 1200μl (10% w/v) and homogenized at 4.0m/s for 45s in a MP Fast-prep 24 5G instrument. After homogenization, the supernatant was divided in half and transferred to two new microcentrifuge tubes. Triton X-100 was added to one of the transferred supernatants to a final concentration of 0.5% and incubated at room temp for 30 minutes to inactivate maSCV2. Infectious and inactivated homogenates were stored at −80°C. Infectious virus titers in tissue homogenate or sera were determined by TCID_50_ assay. Tissue homogenates or sera were serially diluted in infection media in sextuplicate into 96-well plates confluent with Vero-E6-TMPRSS2 cells, incubated at 37°C for six days. After incubation, 10% neutral buffered formalin was added to all wells to fix cells prior to staining and left overnight. Formalin was discarded and the plates were stained with naphthol blue black stain for visualization. Infectious virus titers were determined via the Reed and Muench method. Viral RNA copy number was determined by quantitative polymerase chain reaction (qPCR). A 200 µL aliquot of tissue homogenate or serum was mixed with 1 mL of TRIzol reagent (Invitrogen, Cat#15596026) for RNA extraction. To this, 200 µL of chloroform (Fisher Scientific, Cat#C298-500) was added, followed by centrifugation at 12,000 x g for 15 minutes at 4°C. The clear supernatant was collected and an equal volume of 100% isopropyl alcohol (Fisher Scientific, Cat#A416) was added. This mixture was centrifuged at 12,000 x g for 10 minutes at 4°C. The resulting RNA pellet was washed with 75% ethanol (Fisher Scientific, Cat#BP2818-500), air-dried, and resuspended in 50 µL of nuclease-free water. The RT-qPCR for SARS-CoV-2 N1 gene detection was carried out by adding 2.5 µL of the isolated RNA into a master mix composed of 2.5 µL TaqPath™ 1-Step Multiplex Master Mix (Applied Biosystems, Cat#A28526), 0.75 µL of N1 SARS-CoV-2 RUO qPCR Primer & Probe Kit (IDT, Cat#10006713), and 4.25 µL of nuclease-free water. This mix was added to each well of a MicroAmp™ Optical 384-Well Reaction Plate (Applied Biosystems, Cat#4309849). Serial dilutions of N1 were prepared in 10-fold increments for absolute quantification of copy number. Each sample and standard were run in duplicate. The QuantStudio 12K Flex Real-Time PCR System (Applied Biosystems) was used for amplification, and data analysis was performed using the Design & Analysis Software 2.6.0 to identify SARS-CoV-2 N1.

### 2.10 Placental histology and immunohistochemistry

Placentas were fixed for 72 hours at 4°C in 4% PFA in the ABSL3. Placentas were washed five times with PBS and immersed in 30% sucrose until saturation. Using a Leica CM1950 cryostat, the specimens were cut at 20-μm thickness and mounted on positively charged slides (Fisher Scientific #12-550-15). Routine H&E staining was performed to evaluate the morphological change of the placentas. Within H&E-stained sections, mononucleated trophoblast giant cells, distinguished by their large size and the presence of a single condensed dark blue-purple stained nucleus, were identified and counted under a magnification of 20x. For each placenta, six random images in the labyrinth at the middle level (thickest) of placenta were taken and the count was averaged. For immunohistochemical staining, slides were washed with PBS, which was followed by permeabilization in PBS solution containing 0.05% Triton X-100 and 10% normal goat serum (Invitrogen #50197Z) for 30 min. Placentas were incubated with rabbit anti-vimentin (1:200, Abcam # ab92547), or rabbit anti-cytokeratin (1:200, Dako #Z0622) overnight at 4°C. The next day, sections were rinsed with PBS and then incubated with donkey anti-rabbit (ThermoFisher #R37119) fluorescent secondary antibodies (ThermoFisher #R37115) diluted 1:500 for 3 h at room temperature. DAPI (Roche #10236276001) was applied for counterstaining, followed by mounting with Fluoromount-G (eBioscience #00-4958-02). Images were taken using a Zeiss Axioplan 2 microscope (Jena, Germany) under 5x or 20x magnification. Cell density of vimentin and cytokeratin positive cell quantification was performed using Image J (1.47v). The 20x images were captured from the same batch of experiments, utilizing identical imaging parameters, including exposure time for quantification. After setting the appropriate scale and threshold for positive expression, the percentage of positive expression relative to the entire area was calculated. For each placenta, six random images in the labyrinth at the middle level (thickest) of placenta were taken, and the average fluorescent area calculated for that placenta. One placenta per dam was used and 4–5 dams per group were analyzed.

### 2.11 Cortical thickness measurement

A subset of offspring was randomly selected to be euthanized via decapitation at PND 0 and heads were fixed for 72 hours at 4°C in 4% PFA in the ABSL3. Fetal heads were washed five times with PBS and immersed in 30% sucrose until saturation. Using a Leica CM1950 cryostat, the specimens were cut at 20-μm thickness and mounted on positively charged slides (Fisher Scientific #12-550-15). Nissl staining was performed, and images were taken under × 5 magnification using a Canon EOS Rebel (Tokyo, Japan). Coronal cortical thickness was measured from five random sections at the striatum level of each neonatal brain, as previously described (57). Cortical thickness was measured from both brain hemispheres in each section using ImageJ software, and the average of 10 measurements per specimen was presented. Quantification shown represents the average measurement from a single randomly chosen pup for each dam.(57)

### 2.12 Interferon β and Interleukin 1β ELISA

Interferon β in inactivated right cranial lung or placental homogenate was measured by ELISA according to the manufacturer’s protocol (PBL Assay Science # 42410-1). Interleukin-1β in inactivated placental homogenate was measured by ELISA according to the manufacturer’s protocol (Abcam #100704).

### 2.13 Western Blot

Flash frozen median liver lobes were homogenized in 1X Cell lysis Buffer (Cell Signaling Technology #9803) with 1X Protease Inhibitor cocktail (Sigma-Aldrich #P8340) and sodium fluoride (Fisher Scientific #S299 100) (20μl lysis buffer per mg tissue). Protein lysates were stored at −80°C until analysis. Protein concentration of each lysate was measured using the Pierce BCA Protein Assay Kit (Thermo Fisher Scientific #23225). For each sample, 20μg of protein was subjected to sodium dodecyl sulfate-polyacrylamide gel electrophoresis (SDS-PAGE) on NuPAGE 4-12% Bis-Tris gels (Thermo Fisher Scientific #NP0329). The gel was blotted onto Immobilon-FL PVDF Membrane (Millipore #IPFL00010) and the membranes were blocked with a 1:1 mixture of 1XPBS/Tween-20 solution (Sigma-Aldrich #P3563) and Intercept blocking buffer (LI-COR Biosciences #927-70001) for 30 minutes at room temperature. Membranes were treated with a primary antibody diluted in blocking solution at 4°C overnight on a rocker. Membranes were then washed with PBS-Tween three times and incubated in secondary antibody solutions for one hour at room temperature on a rocker. Membranes were washed three times in PBS-Tween and then imaged on a ProteinSimple FluoroChem Q imager. Individual bands were quantified using Image Studio software (LI-COR Biosciences; version 3.1.4) The signal from each band was normalized against the GAPDH signal and graphed as arbitrary units. Primary antibodies used were rabbit anti-P450 3A4/CYP3A4 (abcam #ab3572) and mouse anti-GAPDH (abcam #ab82450). Secondary antibodies included goat anti-mouse Alexa Fluor 488 (Thermo Fisher Scientific #A11001) and donkey anti-rabbit Alexa Fluor Plus 647 (Thermo Fisher Scientific #A32795).

### 2.14 Viral RNA extraction, sequencing, and analysis

For each sample, 200μl of right cranial lung or nasal turbinate homogenate was mixed with 1mL of TRIzol (Invitrogen #15596026), followed by 200μl of chloroform (Fisher Scientific #C298-500) to extract RNA, and centrifuged at 12,000 g for 15 minutes at 4°C. The clear portion of the supernatant was then pelleted at 12,000 g for 10 minutes along with 500μl of 100% isopropyl alcohol (Fisher #A416) at 4°C. Pelleted RNA was then washed with 75% of ethanol (Fisher Scientific #BP2818-500), air dried and resuspended in 20μl of nuclease-free water. Reverse transcription was carried out using ProtoScript® II First Strand cDNA Synthesis Kit (New England Biolabs #E6560S) with random hexamer mix. The Mpro region was then amplified using forward primer 5’ ACAAAGATAGCACTTAAGGGTGG 3’ and reverse primer 5’ GCGAGCTCTATTCTTTGCACTAA 3’ and Oxford Nanopore sequenced by Plasmidsaurus (SNPsaurus LLC). Consensus sequences for lung and turbinate virus isolates were imported and aligned to Mpro ORF (NC_045512.2) using ClustalO v1.2.3 in Geneious Prime v2023.0.4. Alignments were imported into R v4.1.1, visualized, and annotated using seqvisR v0.2.5. ORF1a and nonstructural protein annotation was visualized using BioRender. Raw FASTQ files for Mpro sequencing has been deposited through SRA under BioProject PRJNA940500 at accession numbers SRR23689223 - SRR23689259.

### 2.15 Statistical analyses

Post infection body mass changes were plotted and the area under the curve (AUC) was calculated to provide individual data points that captured change over time, with AUCs compared with either two tailed unpaired t-test or two-way analysis of variance (ANOVA) followed by *post-hoc* Bonferroni multiple comparisons tests. To compare body mass changes across gestational ages, individual AUCs were subtracted from the average AUC of mock mice at the same gestational age, with the difference from mock AUC compared with two-way ANOVAs followed by *post-hoc* Bonferroni multiple comparisons test. Cumulative clinical scores were analyzed using the Kruskal-Wallis test. Viral titers in lungs from infected dams were analyzed using one-way or two-way ANOVAs followed by *post-hoc* Bonferroni multiple comparisons test. Western blot quantification, IHC quantification, and cortical thickness measurements were analyzed with two-tailed unpaired t tests. Cumulative inflammation scoring, DFCO, IFN-β, and fetal measurements were analyzed with two-way ANOVAs followed by *post-hoc* Bonferroni multiple comparisons test. Pup neurodevelopment results were analyzed with two-way or three-way ANOVAs followed by *post-hoc* Bonferroni multiple comparisons test. Fetal viability data were analyzed with a χ2 test. Data are presented as mean ± SEM or as the median (cumulative clinical score). Mean or median differences were considered statistically significant at *p* < 0.05. Statistical analyses were performed using GraphPad Prism v9.5 (GraphPad Software).

### 2.16 Data availability

Raw FASTQ files for Mpro sequencing has been deposited through SRA under BioProject PRJNA940500 at accession numbers SRR23689223 - SRR23689259. Other data supporting the conclusions of this article are available in the supporting data values.

### 2.17 Study approval

All animal procedures were approved by the Johns Hopkins University Animal Care and Use Committee (MO21H246). SARS-CoV-2 was handled in a BSL-3 containment facility using institution approved biosafety protocols (P2003120104).

## 3 Results

### 3.1 Mouse adapted SARS-CoV-2 causes morbidity in pregnant mice, which increases with gestational age

To evaluate if SARS-CoV-2 caused greater disease in pregnant than nonpregnant mice and if maternal morbidity was impacted by gestational age, we intranasally inoculated outbred pregnant CD1 dams at E6, E10, or E16, roughly corresponding developmentally to human first, second, or third trimesters, respectively (61), or age-matched nonpregnant females with maSCV2(44) or media and measured body mass change as a measure of morbidity. Nonpregnant females (**Figure 1A**) and dams infected at E6 (**Figure 1B**) experienced mild morbidity, losing significant body mass [i.e., approximately 10% of their initial body mass by 4 days post infection (dpi)], but then appearing indistinguishable from mock-inoculated females by 7 dpi. In contrast, dams infected at E10 (**Figure 1C**) or E16 (**Figure 1D**) experienced prolonged maternal morbidity, with E10-infected dams gaining less body mass for the remainder of gestation than mock inoculated dams (**Figure 1C**), and E16-infected dams failing to regain body mass for the remainder of gestation or during lactation as compared to mock-inoculated dams (**Figure 1D**). No mortality was observed in any group. To compare the impact of gestation on maternal morbidity, the change in body mass relative to gestational-age matched mock-inoculated animals (**Figure 1E**) and cumulative clinical scores of disease (**Figure 1F**) were analyzed. maSCV2 infection of pregnant dams at E16 resulted in greater body mass loss and clinical disease than infection of either nonpregnant females or dams at either E6 or E10. Pregnancy and gestational age increase the severity of SARS-CoV-2 outcomes in mice, consistent with human COVID-19 data (1, 10).

**Figure 1.**
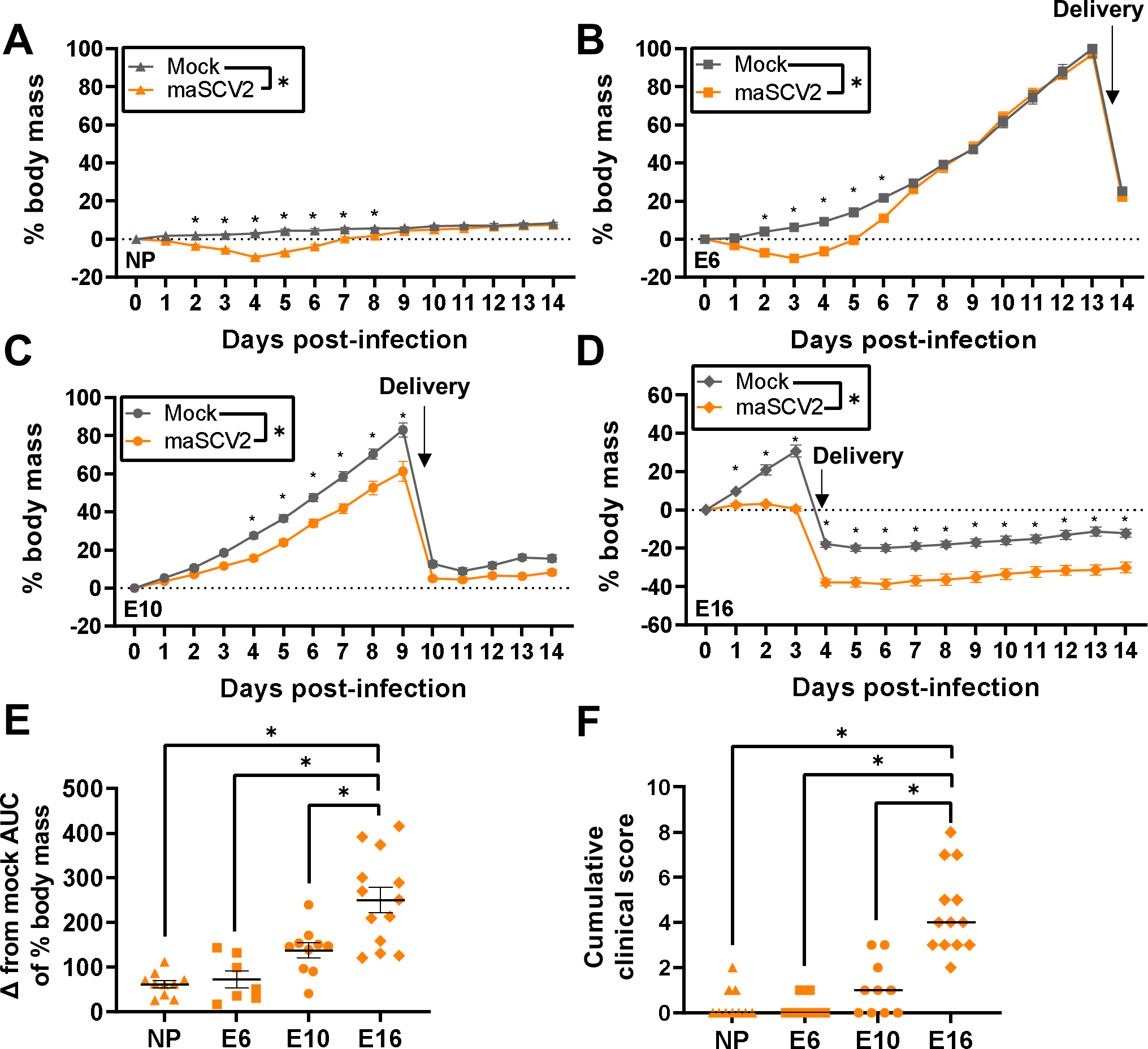
maSCV2 infection of pregnant dams results in gestation-dependent morbidity. Nonpregnant adult females (A) or dams at embryonic day (E) 6 (B), E10 (C), and E16 (D) were intranasally inoculated with 10^5^ TCID_50_ of a mouse adapted SARS-CoV-2 (maSCV2) or mock inoculated with media. Following infection, mice were monitored for change in body mass and clinical signs of disease over fourteen days (A-D). Area under the curve (AUC) of body mass change curves for infected and uninfected animals were calculated, and then the AUC of infected animals was subtracted from the average AUC of mock animals of the same reproductive status and gestational age (E). Clinical scores given to animals included dyspnea, piloerection, hunched posture, and absence of an escape response and are quantified on a score of 0-4. The cumulative clinical score over the 14-day monitoring period is reported for each animal (F). Individual shapes (A-D) or bars (E-F) represent the mean (A-E) or median (F) ± standard error of the mean (A-E) from two independent replications (n =7-13/group) with individual mice indicated by shapes (E-F). Significant differences (*p* < 0.05) were determined by two-way repeated measures ANOVA with Bonferroni post hoc test (A-F, to compare individual timepoints), two tailed unpaired t-test of AUCs (A-F, to compare across all timepoints), one-way ANOVA with Bonferroni post hoc test (E), or Kruskal-Wallis test (F) and are indicated by an asterisk (*).

### 3.2 Pregnant dams infected late in gestation have reduced IFN-β responses, increased viral load, and reduced pulmonary function after infection

Deficits in type 1 IFN signaling are associated with severe COVID-19 in nonpregnant people (62) and mice (63). Pregnancy is associated with downregulation of prototypical cytolytic and anti-viral pathways, including type I IFNs, and upregulation of anti-inflammatory pathways toward mid to late gestation (64). We hypothesized that E16-infected dams would have a reduced type I IFN response after SARS-CoV-2 infection compared to E6-, and E10-infected dams and nonpregnant females. To test this, we infected pregnant dams at E6, E10, and E16 as well as age-matched nonpregnant females with maSCV2 or media and collected lungs at 3 dpi. Infected nonpregnant females as well as E6- and E10-infected dams had significantly increased concentrations of pulmonary IFN-β as compared with matched mock-inoculated females (**Figure 2A**). In contrast, maSCV2 infection at E16 resulted in pulmonary concentrations of IFN-β that were indistinguishable from vehicle-inoculated dams and suppressed relative to either non-pregnant females, E6 dams, or E10 dams infected with maSCV2 (**Figure 2A**). To determine if reduced anti-viral IFN-β concentrations were associated with greater pulmonary virus replication, at 3 dpi, we evaluated infectious viral titers (**Figure 2B**) and viral N1 gene copy numbers (**Supplemental Table 1**) in the lungs of nonpregnant, E6, E10, and E16 pregnant females that were maSCV2-infected. E16-infected dams had significantly greater pulmonary titers of infectious virus and viral RNA than either E6-infected dams, E10-infected dams, or infected nonpregnant females (**Figure 2B, Supplemental Table 1**).

**Figure 2.**
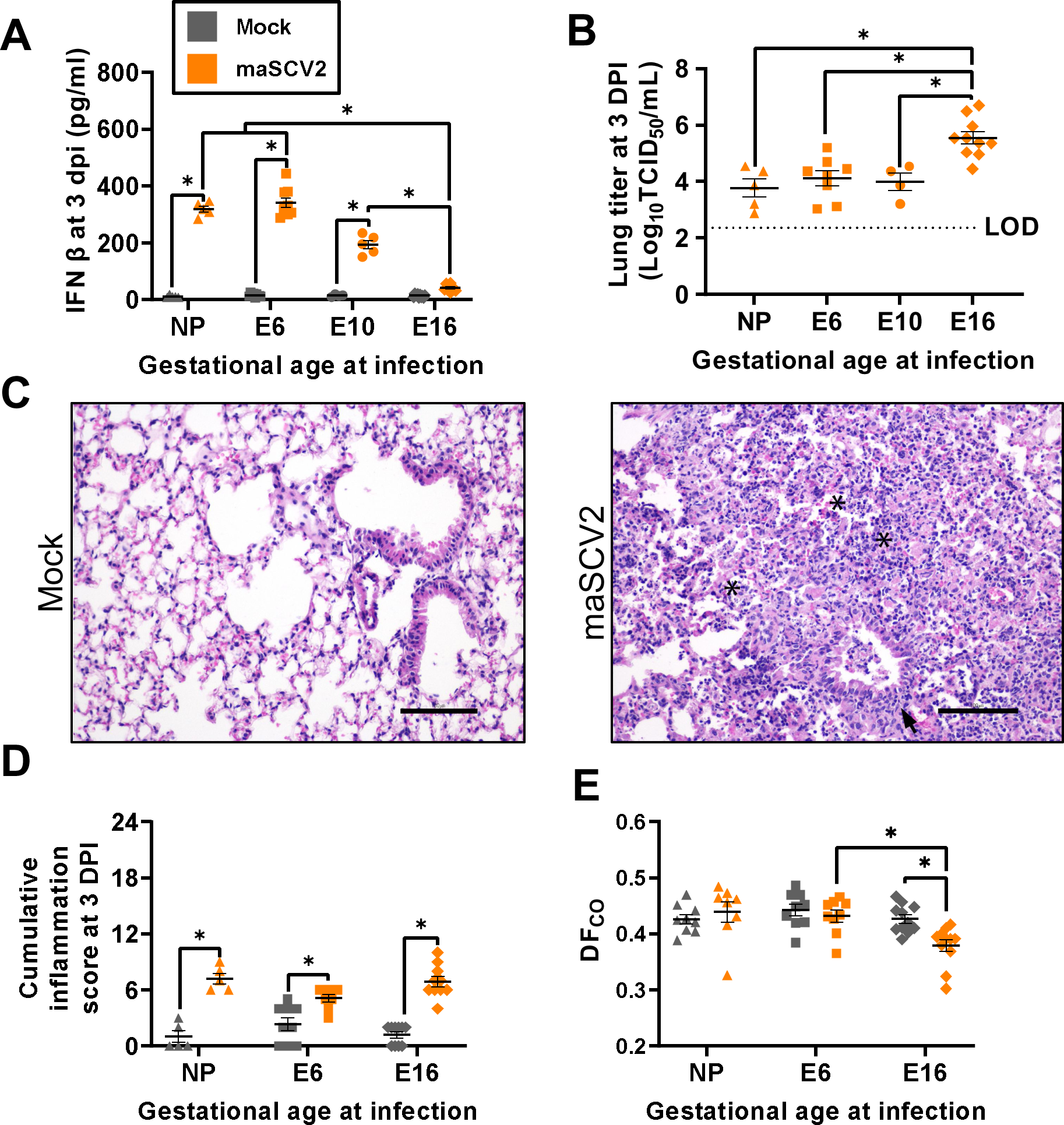
Pregnant dams infected during the third trimester-equivalent have reduced IFN-β responses, increased viral load, and reduced pulmonary function after infection. Nonpregnant adult females or dams at embryonic day (E) 6, E10, and E16 were intranasally inoculated with 10^5^ TCID_50_ maSCV2 or mock inoculated with media and euthanized at three days post infection (DPI) to collect maternal and fetal tissues. IFNβ and viral titers in the right cranial lungs were measured using ELISA (A) and TCID_50_ assay (B), respectively. Sections of fixed left lungs were stained by hematoxylin & eosin (H&E) to evaluate lung inflammation and images were taken at 20x magnification, with representative images of lungs maSCV2 or mock-inoculated at E16 shown (C). Asterisks (*) indicate intra-alveolar necrosis and inflammatory infiltrates, and arrows indicate peribronchiolar inflammatory infiltrates. Histopathological scoring was performed by a blinded board-certified veterinary pathologist to measure cumulative inflammation scores (D). A subset of mice were tracheostomized at three DPI to measure pulmonary function through the diffusion capacity for carbon monoxide (DF_CO_) prior to euthanasia (E). Bars represent the mean (A-E) ± standard error of the mean from at least two independent replications (n =4-11/group) with individual mice indicated by shapes. Significant differences (*p* < 0.05) were determined by two-way ANOVA with Bonferroni post hoc test (A,D,E) or one-way ANOVA with Bonferroni post hoc test (B) and are indicated by an asterisk (*). Scale bar: 100 µm. LOD indicates the limit of detection.

To determine if greater viral replication contributed to worse pulmonary outcomes, at 3dpi, we evaluated pulmonary histopathology (**Figure 2C-D**, **Supplemental Figure 1**), and diffusion capacity (DF_CO_; **Figure 2E**) in the lungs of nonpregnant, E6, or E16 pregnant females that were either maSCV2 or mock-infected. maSCV2 infection induced pulmonary histopathological changes, including intra-alveolar necrosis and inflammatory cell debris, and peribronchiolar, and perivascular mononuclear inflammatory infiltrates that were observed in nonpregnant (**Supplementary Figure 1A**, representative image), E6- (**Supplemental Figure 1B**, representative image), and E16- (**Figure 2C**, representative image) infected mice (**Figure 2D**, scoring) to equivalent levels. maSCV2 infection at E16 significantly reduced pulmonary function, as measured by DF_CO_, which was not observed following maSCV2 infection at E6 or in nonpregnant females. These data suggest that late gestation is associated with reduced antiviral responses, greater virus replication, and reduced pulmonary function.

### 3.3 SARS-CoV-2 infection late in gestation disrupts trophoblasts and cytokine concentrations in the placenta

As placental pathology has been observed during COVID-19 (65–67), we next investigated if maSCV2 could infect or cause damage to the placenta. Dams were mock- or maSCV2-infected at E10 [when the placenta is formed (61)] or E16 and euthanized at 3 dpi. Placentas, fetal tissues, and maternal sera were analyzed for infectious virus and viral RNA (**Supplemental Table 1**), with placentas further analyzed for tissue damage (**Figure 3**). All placentas, fetal tissues, and maternal sera were negative for viral RNA and infectious virus (**Supplemental Table 1**), consistent with human reports that direct placental infection and vertical transmission during COVID-19 is rare (23, 24). Despite no detectable infectious virus or viral RNA, placentas from E16-infected dams had reduced numbers of mononuclear trophoblast giant cells (**Figure 3A** for representative images, and **Figure 3B** for quantification), suggestive of damage to the trophoblast-endothelial cell barrier, which separates maternal and fetal blood in the labyrinth of the murine placenta (68). Staining for cytokeratin (trophoblasts, **Figure 3C, E)** and **Supplemental Figure 2A**) and vimentin (endothelial cells, **Figure 3D, F** and **Supplemental Figure 2B**) was performed and revealed a significant loss of trophoblasts, but not endothelial cells, in placentas from E16-infected compared with mock-infected dams (**Figure 3C-F-C**). These data illustrate disruption of the maternal and fetal barrier in the absence of direct viral infection or vertical transmission. Moreover, cell numbers in placentas of E10-infected dams did not differ from placentas of mock-inoculated dams (**Supplemental Figure 2A-B**). These data suggest that placental damage may be associated with the more severe maternal disease seen with infection at E16, potentially due to maternal immune activation or sickness behavior (69–71), which will require further studies for elucidation.

**Figure 3.**
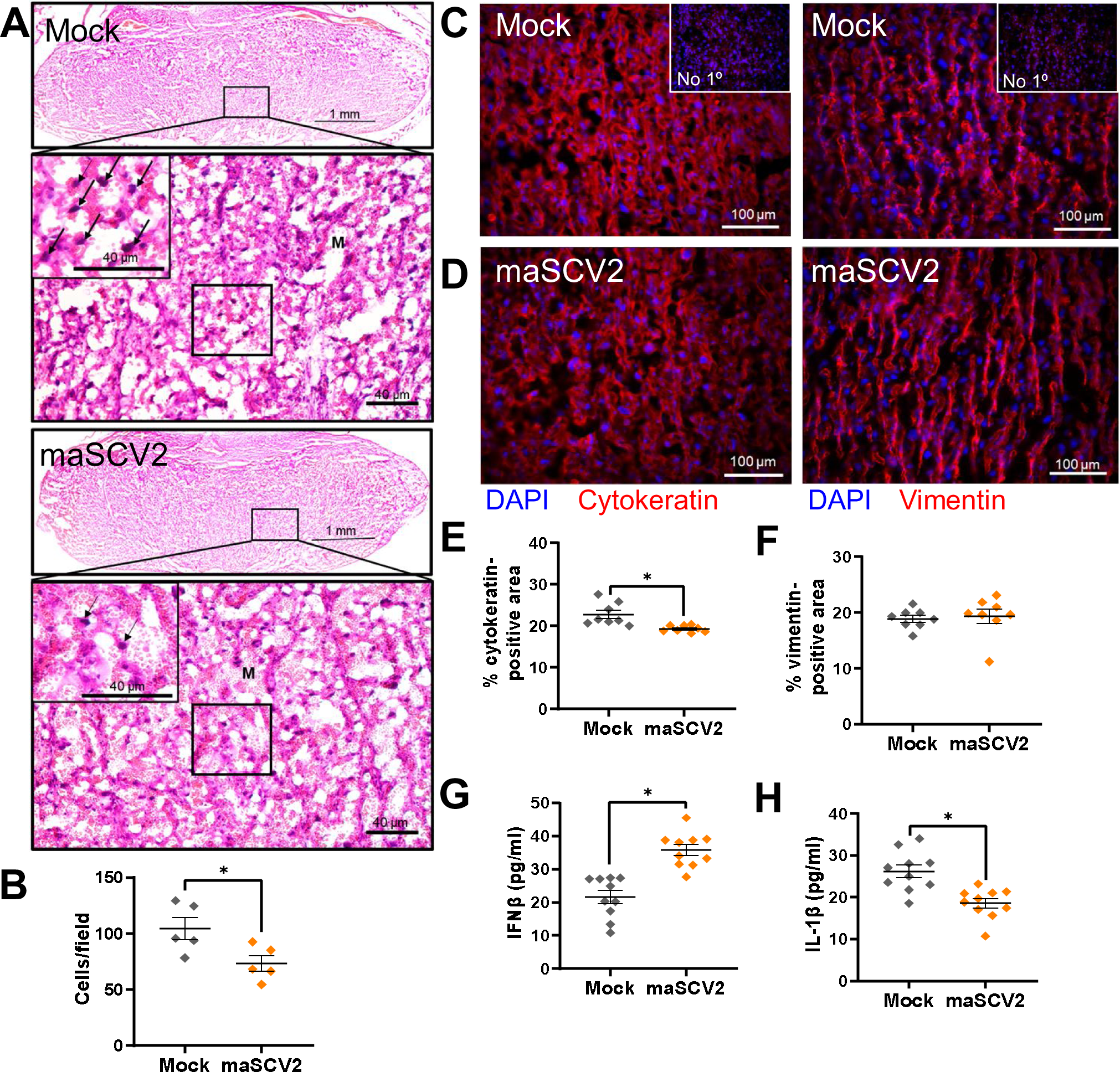
Third trimester-equivalent maSCV2 infection disrupts the trophoblast layer of the placental labyrinth zone and cytokine concentrations. At embryonic day (E) 16, pregnant dams were intranasally inoculated with 10^5^ TCID_50_ of maSCV2 or mock inoculated with media and euthanized at 3 dpi to collect placentas. Representative H&E images (A) were taken at 5x (upper panels) and 20x magnification (lower panels, and specific areas of interest further zoomed 1.75 fold (black box). Within H&E-stained placentas, arrows indicate trophoblast giant cells and Ms indicate maternal blood spaces. Mononucleated trophoblast giant cells were identified and counted at 20x magnification (B). Placentas were immunostained for cytokeratin (C, red) to mark trophoblasts or vimentin (D, red) to mark endothelial cells and DAPI (blue) to label nuclei, with controls without primary antibody run in parallel. Representative images were taken at 20x magnification. Quantification of the percentage positive area for each marker is shown (E-F). Placentas were homogenized and analyzed by ELISA for IFN-β (G) IL-1β (H). Bars represent the mean ± standard error of the mean (n =5-10/group) with each shape indicating 1 placenta and, for analysis of images is the mean quantification or count of 6 fields of view. Significant differences (*p* < 0.05) were determined by unpaired two tailed t-test and are indicated by an asterisk (*). Scale bar: 1mm (A, upper panels/group), 40 µM (A, lower panels/group), or 100 µm (C-D)

Altered concentrations of cytokines, including IFN-β and IL-1β, in the placenta are associated with placental damage (56, 69, 72, 73). As such, we measured IFNβ and IL-1β in placentas of dams that were maSCV2 infected or mock inoculated at E10 (**Supplemental Figure 2C-D**) or E16 (**Figure 3G-H**). Maternal maSCV2 infection at E16, but not E10, resulted in increased concentrations of IFN-β and reduced concentrations of IL-1β in the placenta relative to mock-inoculated dams (**Figure 3G-H**, **Supplemental Figure 2C-D**) These data suggest that maSCV2 infection shifted the balance of these two counter regulatory cytokines in the placenta (74, 75), with placental IFN-β and IL-1β concentrations being correlated, regardless of infection status or timing of infection (**Supplemental Figure 2E**).

### 3.4 SARS-CoV-2 infection late in gestation causes intrauterine growth restriction

COVID-19 during human pregnancy is associated with adverse pregnancy and fetal outcomes including preterm birth, stillbirth, small size for gestational age, and reduced birth weight (15). To evaluate if the maSCV2-induced maternal morbidity and placental damage observed after infection at E16 was associated with adverse pregnancy or fetal outcomes, we inoculated dams with maSCV2 or media at E6, E10, or E16, with a subset of dams euthanized at 3 dpi to evaluate fetal viability and the remainder followed to evaluate birth outcomes. Neither fetal viability (**Figure 4A**) nor litter size (**Figure 4B**) was affected by maSCV2 infection during pregnancy at any gestational age. maSCV2 infection at E6 or E10 did not result in reductions in fetal growth relative to fetuses from mock-inoculated dams (**Figure 4C-E**). In contrast, maSCV2 infection at E16 led to significantly smaller pups in terms of mass, length, and head size relative to fetuses from mock-infected dams (**Figure 4C-E**). Collectively, fetuses from E16-infected dams had greater growth restriction than fetuses from either E6- or E10-infected dams (**Figure 4C-E**). Reduced birth size was not mediated by pre-term birth as all dams, regardless of infection, delivered at approximately E20 (76). These data indicate that maSCV2 infection during the third trimester-equivalent of pregnancy results in intrauterine growth restriction, which was not observed when infection occurs earlier during gestation.

**Figure 4.**
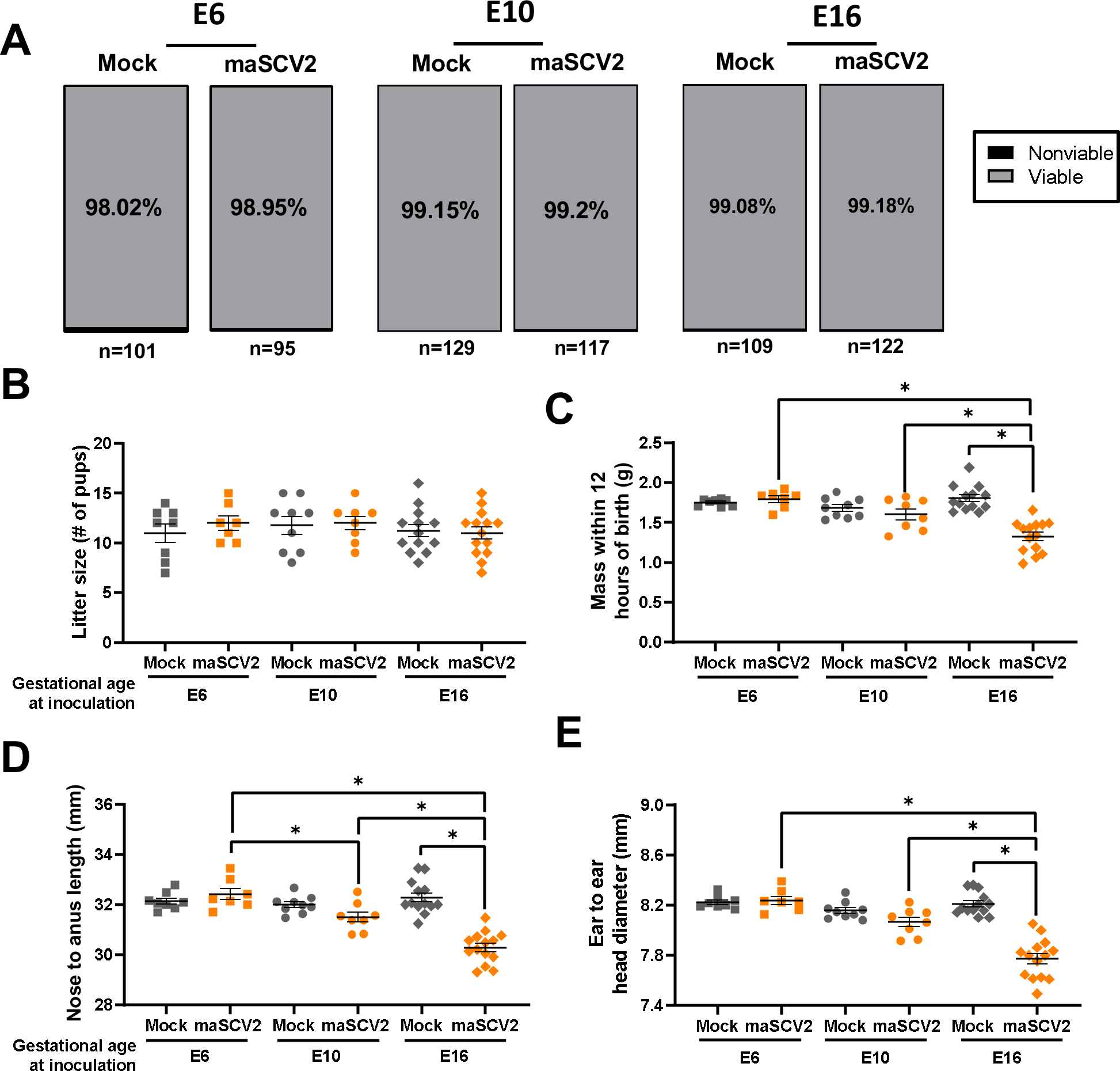
Third trimester-equivalent maSCV2 infection causes intrauterine growth restriction. At embryonic day (E) 6, E10, or E16, pregnant dams were intranasally inoculated with 10^5^ TCID_50_ of maSCV2 or mock inoculated with media. At 3 dpi, a subset of dams were euthanized, and fetal viability was determined as the percentage of fetuses within the uterus (A, n= total number of fetuses from 8-12 dams per group from two independent replicates). Fetuses were counted as nonviable if they were smaller or discolored compared to gestational age-matched live fetuses or if a fetus was absent at an implantation site. A subset of dams were followed into the postnatal period to characterize adverse birth outcomes. At postnatal day 0 (PND0) overall litter size (B), pup mass (C), pup body length (D), and pup head diameter (E) were measured. Average measurements of each independent litter were graphed to account for litter effects (B-E). Bars represent the mean ± standard error of the mean from two independent replicates (n =7-14/group) with the average of individual litters indicated by shapes. Significant differences (*p* < 0.05) were determined by χ2 (A) or two-way ANOVA with Bonferroni post hoc test (B-E) and are indicated by an asterisk (*).

### 3.5 Offspring of SARS-CoV-2 infection late in gestation display cortical thinning and reduced neurodevelopmental behaviors

In addition to adverse perinatal outcomes, COVID-19 during pregnancy also has been associated with an increased risk of neurodevelopmental disorders in infants within their first year of life (20, 21). As such, we evaluated offspring of mock-inoculated or maSCV2-infected dams at E16 for reduced cortical thickness at postnatal day (PND) 0 and delayed neurobehavioral function at PND 5. Offspring of E16-infected dams had significant cortical thinning in comparison to offspring from mock-inoculated dams (**Figure 5A-B**), consistent with their reduced head diameter (**Figure 4E**). Offspring of E16-infected dams displayed delayed surface righting (**Figure 4C**), cliff aversion (**Figure 4D**), and negative geotaxis (**Figure 4E**) as compared with offspring from mock-infected dams. Male offspring were more affected by maternal infection at E16 than female offspring, consistent with literature indicating that males are more severely impacted by *in utero* insults (77, 78), including SARS-CoV-2 infection (21). Offspring of dams that were either maSCV2- or mock-infected at E6 or E10 also were subjected to neurobehavioral testing, and no effect of either maternal infection or sex of offspring was observed (**Supplemental Figure 3**). These data highlight that infection with maSCV2 during the third trimester-equivalent of pregnancy causes both short and long-term adverse fetal outcomes, in the absence of vertical transmission and consistent with human literature (2, 20, 21).

**Figure 5.**
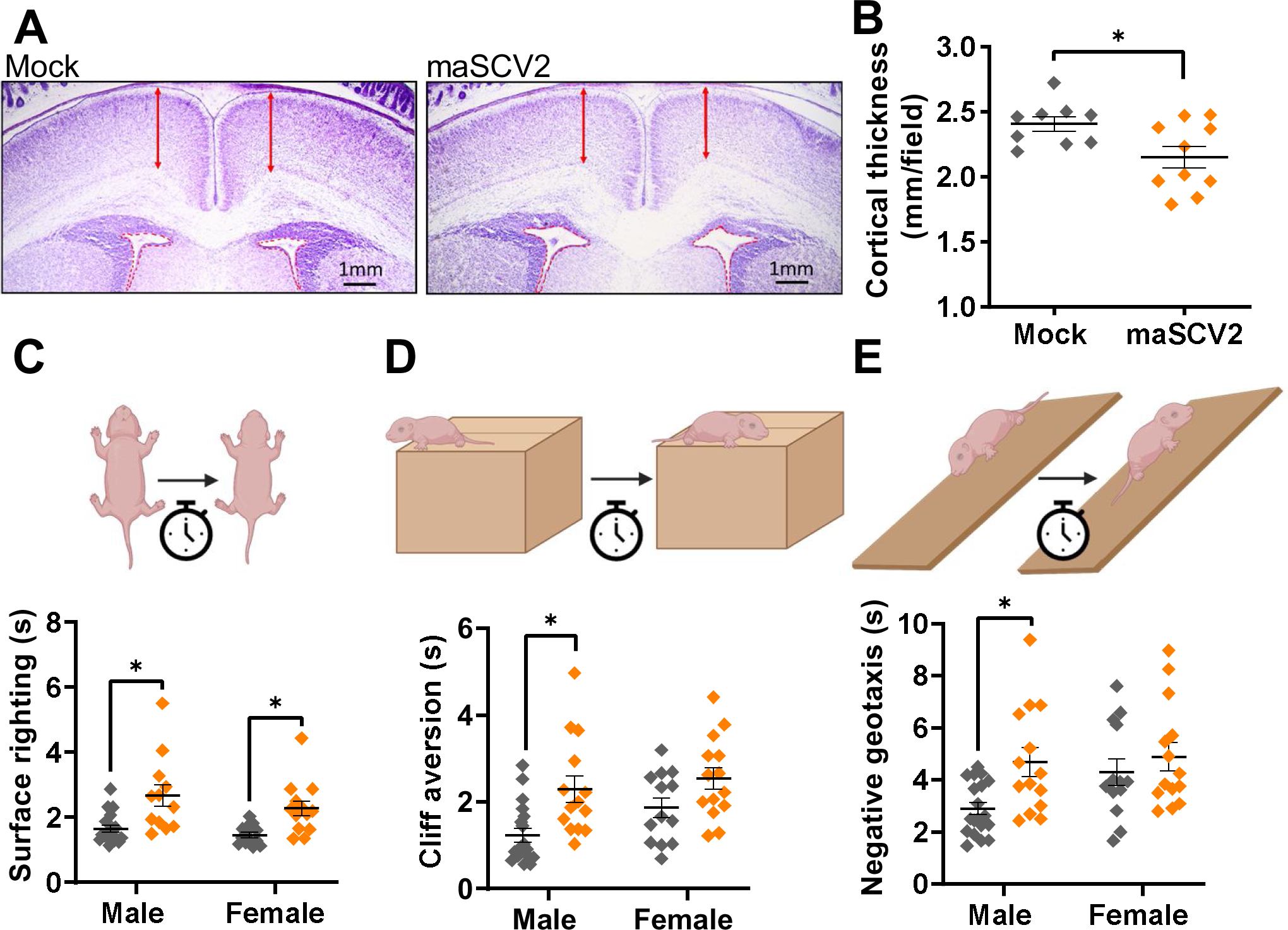
Offspring of dams infected with maSCV2 during the third trimester-equivalent display cortical thinning and reduced neurodevelopmental function. At embryonic day (E) 16, pregnant dams were intranasally inoculated with 10^5^ TCID_50_ of maSCV2 or mock inoculated with media. At PND0, a randomly selected subset of pups were euthanized via decapitation to collect fetal heads, which were fixed, sliced, and Nissl stained. Cortical thickness (A, red arrows) was measured from both brain hemispheres per pup and quantified as the average of 10 measurements per pup, with a single pup randomly chosen per dam (B, n=9-10 independent litters/group from 2 independent replicates). A subset of offspring were followed to PND5, sexed, and the neurobehavioral assays of surface righting (C), cliff aversion (D), and negative geotaxis (E) were performed to measure neurological development. 1-2 pups per sex per dam were subjected to each test subsequently, with 3 trials given per test, and each pup’s best trial for each test was reported (C-E, n=9-10 independent litters/group from 2 independent replicates). Bars represent the mean ± standard error of the mean with each shape indicating 1 pup. Significant differences (*p* < 0.05) were determined by unpaired two tailed t-test (B) or two-way ANOVA with Bonferroni post hoc test (C-E) and are indicated by an asterisk (*). Graphics built with Biorender.com.

### 3.6 Ritonavir-boosted nirmatrelvir treatment prevents morbidity and reduces pulmonary viral titers following SARS-CoV-2 infection late in gestation

Because of the increased risk of severe COVID-19 and adverse fetal outcomes, pregnant individuals are recommended to receive the antiviral ritonavir-boosted nirmatrelvir in the United States (33, 34). There is, however, limited data on its efficacy during pregnancy, with human and animal studies primarily focused on evaluating safety and toxicity (38, 42). Additionally, studies evaluating nirmatrelvir’s efficacy in nonpregnant animals utilized high doses of nirmatrelvir alone in lieu of boosting with ritonavir (52, 79). To better reflect the doses administered to pregnant individuals, we first evaluated the efficacy of nirmatrelvir and ritonavir at doses calculated to be the mouse equivalent to a human doses (49) in nonpregnant females compared to high dose nirmatrelvir alone. Mouse equivalent dosing of nirmatrelvir and ritonavir was equivalent to high dose nirmatrelvir alone at preventing maSCV2 induced morbidity (**Supplemental Figure 4A**) and reducing pulmonary viral loads (**Supplemental Figure 4B**) in nonpregnant females. As CYP3A enzymes are responsible for the metabolism of nirmatrelvir (80), we evaluated liver CYP3A in pregnant dams at E16 and age-matched non-pregnant females and found no difference in total expression (**Figure 6A**), further supporting the use of mouse-equivalent doses of nirmatrelvir and ritonavir during pregnancy.

**Figure 6.**
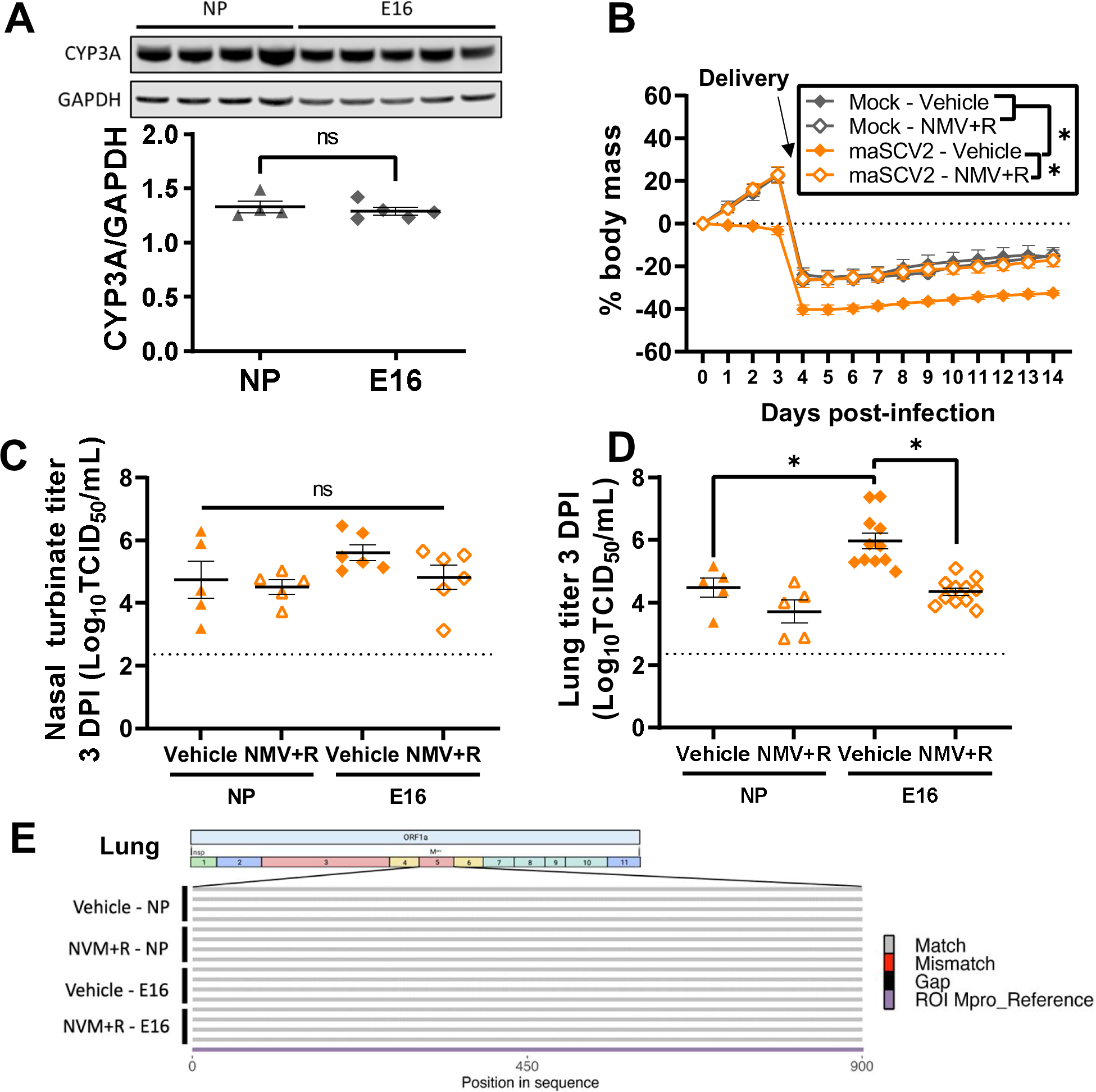
Ritonavir-boosted nirmatrelvir mitigates maternal morbidity and reduced viral titers in the lungs of pregnant dams. Uninfected adult nonpregnant and pregnant (i.e., embryonic day (E)16) females were euthanized, liver tissue collected, and western blots performed to quantify the amount of overall CYP3A expression (A, n=4-5/group). At E16, pregnant dams or age-matched nonpregnant females were intranasally infected with 10^5^ TCID_50_ of maSCV2 or mock inoculated with media. Starting at 4 hours post infection and continuing twice daily for 5 days or until tissue collection, mice were treated with 1.7 mg nirmatrelvir and 0.6 mg ritonavir per dose or vehicle and were monitored for changes in body mass for fourteen days (B, n=6/group from two independent replicates). A subset of dams were euthanized at 3 dpi, and nasal turbinate and lung tissue were collected, and viral titers were measured by TCID_50_ assay (C-D, n=5-11/group). RNA was extracted from lung homogenate, reverse transcribed using ProtoScript® II First Strand cDNA Synthesis Kit, the M^pro^ region amplified, and Oxford Nanopore sequenced by Plasmidaurus. Consensus sequences were imported and aligned to M^pro^ using ClustalO v1.2.3 in Geneious Prime v2023.0.4. Alignments were imported into R v4.1.1., visualized, and annotated using seqvisR v0.2.5 (E, n=4/group). Bars represent the mean ± standard error of the mean from two independent replications with individual mice indicated by shapes (A,C,D). Significant differences (*p* < 0.05) were determined by two tailed unpaired t-test (A), two-way ANOVA with Bonferroni post hoc test of AUCs (B), or two-way ANOVA with Bonferroni post hoc test (C-D) and are indicated by an asterisk (*). Sequence graphic built using Biorender.com. LOD indicates the limit of detection.

To evaluate the efficacy of ritonavir-boosted nirmatrelvir during pregnancy, we treated maSCV2 and mock-infected dams twice daily with mouse-equivalent doses of nirmatrelvir and ritonavir or vehicle for 5 days (50), starting at 4 hours after infection. maSCV2-infected dams treated with vehicle failed to gain mass during the remainder of pregnancy and had reduced mass compared to mock-inoculated dams through lactation (**Figure 6B**). In contrast, treatment of maSCV2-infected dams with ritonavir-boosted nirmatrelvir prevented maternal morbidity and resulted in morbidity curve AUCs that were equivalent to those of mock-inoculated dams (**Figure 6B**). Ritonavir-boosted nirmatrelvir did not significantly reduce infectious viral loads in the nasal turbinates of pregnant or nonpregnant females (**Figure 6C**). In the lungs, however, ritonavir-boosted nirmatrelvir reduced viral loads in pregnant, but not nonpregnant, females compared to vehicle-treated comparators, likely because infected vehicle-treated nonpregnant females already had lower viral loads than infected pregnant vehicle-treated dams (**Figure 6D**). We next determined if treatment with ritonavir-boosted nirmatrelvir selected for mutations in the coding region corresponding to the gene that encodes for the SARS-CoV-2 MPRO protease. The sequences encoding M^PRO^ did not differ between viral RNA obtained from ritonavir-boosted nirmatrelvir treated mice and vehicle treated mice, regardless of either pregnancy status or tissue type (**Figure 6E**, **Supplemental Figure 5**), suggesting that ritonavir-boosted nirmatrelvir is not selecting for mutations that would potentially reduce its efficacy, at least by 3 dpi.

### 3.7 Ritonavir-boosted nirmatrelvir treatment prevents adverse offspring outcomes induced by SARS-CoV-2 infection late in gestation

To evaluate if ritonavir-boosted nirmatrelvir prevented adverse fetal and offspring outcomes, offspring of E16-infected and mock-inoculated dams treated with ritonavir-boosted nirmatrelvir, or vehicle were evaluated at birth and PND5. Offspring of maSCV2-infected dams treated with vehicle were significantly smaller than offspring of mock-inoculated dams in mass (**Figure 7A**), length (**Figure 7B**), and head diameter (**Figure 7C**) at birth and demonstrated significant delays in surface righting (**Figure 7D**), cliff aversion (**Figure 7E**), and negative geotaxis (**Figure 7F**) at PND5, with greater neurobehavioral delays in males than females (**Figure 7D-F**). Offspring of maSCV2-infected dams treated with ritonavir-boosted nirmatrelvir, however, did not differ from offspring of mock-inoculated dams in any size measures at birth (**Figure 7A-C**) or neurobehaviors at PND5 (**Figure 7D-F**). Ritonavir-boosted nirmatrelvir treatment prevented maSCV2-induced intrauterine growth restriction and neurobehavioral deficits in both males and females. Offspring of mock-inoculated dams treated with ritonavir-boosted nirmatrelvir did not differ from offspring of mock-inoculated dams in any offspring measure (**Figure 7A-F**), consistent with reproductive studies in rabbits which did not find toxicity during pregnancy (42) Overall, these findings suggest that treatment with ritonavir-boosted nirmatrelvir during pregnancy can not only reduce maternal pulmonary viral load, but prevents maternal morbidity, and mitigates adverse fetal and offspring outcomes.

**Figure 7.**
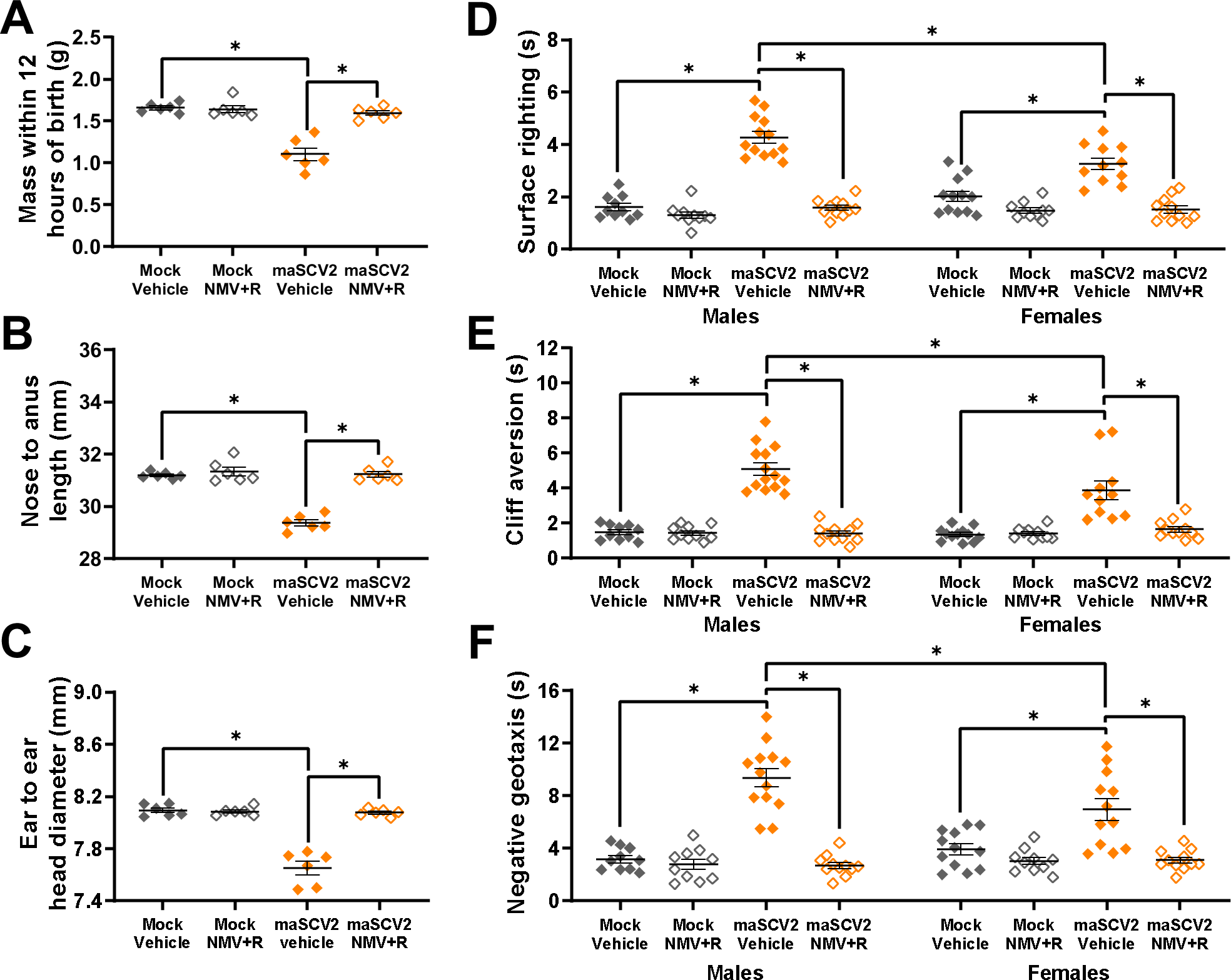
Ritonavir-boosted nirmatrelvir prevents adverse offspring birth outcomes and neurodevelopmental deficits associated with maternal maSCV2 infection. At embryonic day (E)16, pregnant dams were intranasally inoculated with 10^5^ TCID_50_ of maSCV2 or mock inoculated with media. Starting at 4 hours post infection and continuing twice daily for 5 days or until tissue collection, mice were treated with 1.7 mg nirmatrelvir and 0.6 mg ritonavir per dose or vehicle. At PND0, a subset of pups were measured for pup mass (A), pup length (B), and pup head diameter (C). Average measurements of each litter were graphed to account for litter effects (A-C, n=6 independent litters/group from 2 independent replicates). A subset of offspring were followed to PND5, sexed, and the neurobehavioral assays of surface righting (D), cliff aversion (E), and negative geotaxis (F) were performed to measure neurological development. 1-2 pups per sex per dam were subjected to each test subsequently, with 3 trials given per test, and each pup’s best trial for each test was reported (D-F, n=6-8 independent litters/group from 2 independent replicates). Bars represent the mean ± standard error of the mean with each shape indicating 1 litter’s average (A-C) or 1 pup (D-E). Significant differences (*p* < 0.05) were determined by two-way ANOVA with Bonferroni post hoc test (A-C) or three-way ANOVA with Bonferroni post hoc test (D-F) and are indicated by an asterisk (*).

## 4 Discussion

Animal models of COVID-19 are powerful tools to study pathogenesis, consider risk-altering conditions such as pregnancy, and evaluate therapeutic interventions (81, 82). In the current study, we established a mouse model of SARS-CoV-2 infection during pregnancy that recapitulates many of the clinical findings of COVID-19 during human pregnancy. Pregnant dams infected with maSCV2 in late gestation experienced the most severe disease, exhibiting reduced pulmonary function and increased viral titers, while their offspring were small for gestational age and had neurodevelopmental delays. These findings are consistent with observations in humans where pregnant individuals with COVID-19, especially in mid to late gestation, have greater risk of severe disease, resulting in increased hospitalization and critical care admission (10, 11). Virological, biological, and social factors, including SARS-CoV-2 infectious dose and variant, preexisting immunity, and access to healthcare likely contribute to the diversity of adverse fetal outcomes observed with human COVID-19 during pregnancy (15, 18, 32, 83). Our mouse model does not account for all of these factors, which may explain the selective manifestation of adverse fetal outcomes, such as reduced birth mass and neurodevelopmental outcomes, that are worse in male than female offspring (16, 17, 20, 21) Our model did not capture other aspects of COVID-19 during pregnancy, including preterm birth or stillbirth (5, 15, 18, 22), that have been observed in human cases.

At the maternal-fetal interface, intranasal maSCV2 infection resulted in placental alterations without direct virus infection, which is in accordance with the hallmarks of placental damage, inflammation, and maternal immune cell infiltration observed in placentas from mothers with COVID-19 during pregnancy (65–67). After characterizing the negative outcomes of maSCV2 infection in pregnancy, we used our model to assess the efficacy of ritonavir-boosted nirmatrelvir at a mouse-equivalent dose to what pregnant humans receive. This antiviral regimen was well tolerated by pregnant dams, reduced pulmonary virus titers, mitigated maternal morbidity, and prevented adverse offspring outcomes. Observational studies in human pregnancies indicate that ritonavir boosted-nirmatrelvir does not pose safety or toxicity risk to pregnant individuals (38), and may reduce COVID-19 symptoms without requiring additional medical interventions (39).

In addition to recapitulating aspects of human COVID-19 during pregnancy, our model identified reduction in pulmonary IFN-β secretion after infection late in gestation and a corresponding increase in pulmonary viral titer as critical mediators of worse outcomes in late compared with early gestation. As deficits in type 1 IFN signaling have been associated with severe COVID-19 in both nonpregnant individuals (62) and mice (63), our data suggest that maternal morbidity may, in part, be due to an inability of pregnant dams to control viral replication because of a reduced type I IFN responses, particularly during late gestation. This potential mechanism of severe disease is consistent with immunological alterations of mouse and human pregnancy where the maternal immune response shifts to an anti-inflammatory profile to support the semi-allogenic fetus and diverts from anti-viral and cytotoxic activity (12). Moreover, in human pregnancy, there is a documented decline in early anti-viral effector cells and products including natural killer cells and type I IFNs (64).

The adverse maternal and fetal outcomes of SARS-CoV-2 infection during pregnancy are like those observed in other mouse models of viral pathogenesis during pregnancy including Zika virus (ZIKV) and influenza A virus (IAV) infection. Mouse models of ZIKV infection during pregnancy have shown that adverse fetal and neonatal outcomes including congenital abnormalities, reduced cortical thickness, and neurobehavioral deficits (54, 57), are mediated in part by transplacental virus transmission and acute placental inflammation (54, 56). While vertical transmission during ZIKV infection contributes to adverse outcomes, we and others have shown that the maternal immune response, including elevated production of IL-1β, also plays a key role in pathogenesis (54, 84). Vertical transmission of virus during COVID-19 is largely unseen in humans (23, 24, 66), and in mice the placental pathology following maSCV2 infection occurred without vertical transmission. These data further highlight that adverse neonatal outcomes are not exclusive to vertical transmission of viruses, but by maternal immune activation and damage at the maternal-fetal interface. Mouse models of IAV infection during pregnancy further demonstrate maternal morbidity and mortality which is more severe in pregnant than nonpregnant animals (68, 71). Reduced type I IFN responses and greater viral loads in the lungs in pregnant dams late in gestation also have been observed in IAV infection (85), further supporting that pregnancy-associated suppression of type I IFNs is a mechanism of severe maternal disease after respiratory virus infection.

Mouse models of viral infection during pregnancy are a valuable tool to assess the safety and efficacy of therapeutics to prevent adverse maternal and fetal outcomes. Our results support the efficacy of ritonavir-boosted nirmatrelvir for COVID-19 during pregnancy. While human studies of ritonavir-boosted nirmatrelvir during pregnancy are still needed, these findings provide a foundation for future human clinical trial design to include pregnant patients. Current approaches to assessing antiviral therapeutics in preclinical animal models include reproductive toxicity studies using supraphysiological doses but neglect to evaluate if pregnancy alters efficacy (42, 86, 87). Therefore, future preclinical models of antiviral therapies must be designed carefully to consider the complex interactions between pregnancy, viral pathogenesis, and drug pharmacokinetics. Mouse models of ZIKV antiviral treatment during pregnancy have illustrated the ability of maternal antiviral treatment to prevent vertical transmission to fetuses (88), a major adverse outcome associated with ZIKV infection during pregnancy. Pregnant individuals are largely excluded from clinical trials (29), which has contributed to a reduced uptake of antivirals and vaccines in pregnant populations, including COVID-19 therapeutics (89, 90). This exclusion is concerning because pregnant individuals and their neonates are highly vulnerable to many pathogens (91, 92). With further development of mouse models of viral infection and novel therapeutics in pregnancy, however, preclinical studies can guide clinical trial design and promote the inclusion of pregnant populations. By considering pregnancy in clinical trials, access and uptake of protective therapeutics during pregnancy can be improved.

## Supporting information

Supporting Data Values

## 5 Acknowledgements

The authors would like to thank Dr. Ralph Baric as well as the Klein, Pekosz, Davis, and Baumgarth laboratories for discussions about these data, and Ariana Campbell for early assistance with animal studies. We would also like to thank Dr. Jason Villano and the expert animal care staff at the Johns Hopkins School of Medicine for assistance with maintenance of SARS-CoV-2-infected dams.

## 7 Contributions

SK, AP, IB, PC, and JP conceptualized and designed the experiments. PC and JP performed animal experiments. AP, WZ, and RZ grew and quantified viruses and homogenized and inactivated tissue. PC and JP performed ELISA and Western blot. KM imaged and scored lung histology slides. PC, JP, HR, and WM performed DFco analysis. JL and AL stained, imaged, and analyzed placental and fetal head tissue. EA analyzed sequencing data. PC and JP statistically analyzed and graphed data. PC, JP, and SK wrote the manuscript with input from all authors. All authors read and provided edits to drafts and approved the final submission. The order of authors was determined based on contributions to the overall design, experimentation, analyses, and writing.

