## Supplemental Figures for "Adverse outcomes in SARS-CoV-2 infected pregnant mice are gestational age-dependent and resolve with antiviral treatment"

### Supplemental Figure 1.

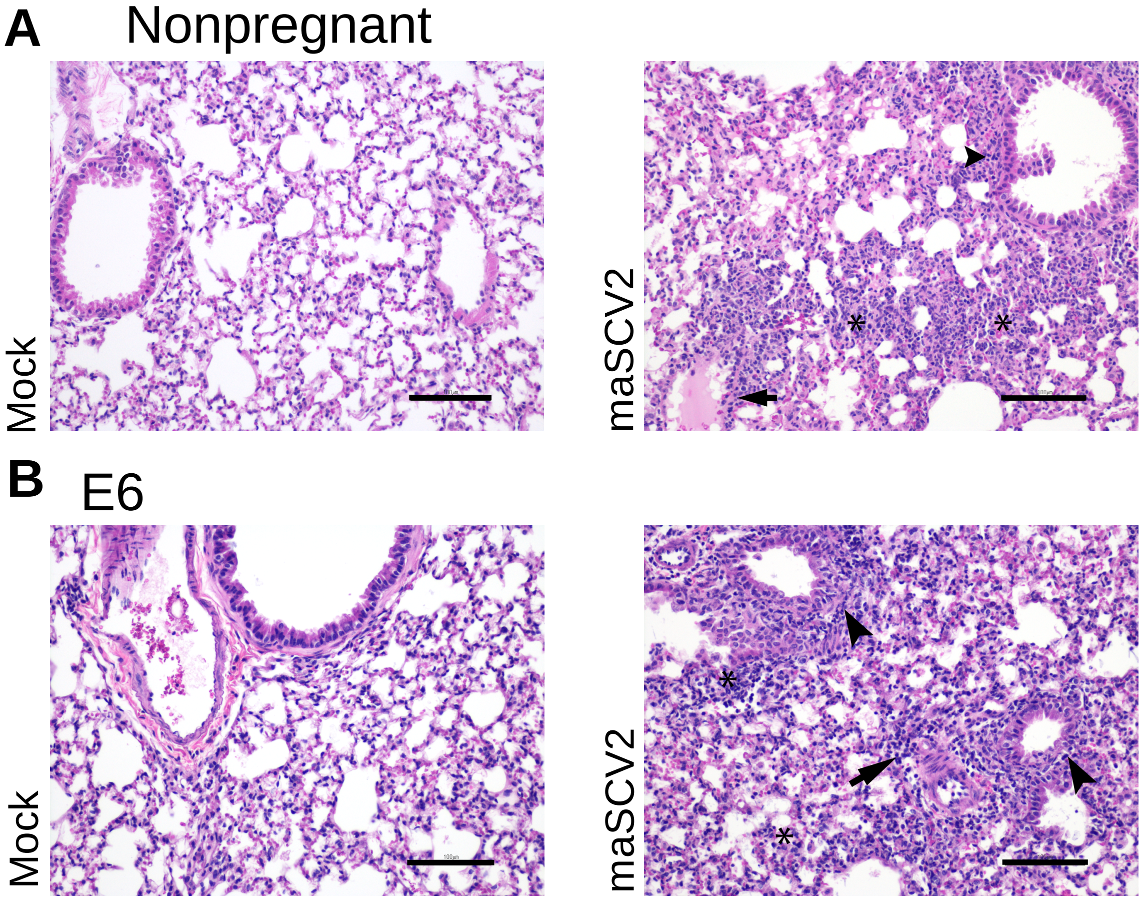


**Supplemental Figure 1. maSCV2 infection induces pulmonary inflammation in nonpregnant female and pregnant first trimester-equivalent mice**
Nonpregnant and pregnant mice at embryonic day (E) 6 that were infected with maSCV2 or mock inoculated with media were euthanized at three DPI to collect maternal and fetal tissues. Left lungs were inflated and fixed using zinc-buffered formalin, sliced into 3-mm blocks, embedded in paraffin, and sectioned to 5μm. Sections were mounted on glass slides and stained with H&E to evaluate lung inflammation and representative images of lungs from nonpregnant female mice (A-B) and pregnant mice at E6 (C-D) were taken at 20x magnification. Intra-alveolar, perivascular and peribronchiolar inflammatory infiltrates are indicated with an asterisk, arrow, and arrowhead, respectively. Scalebar: 100 μm

### Supplemental Figure 2.

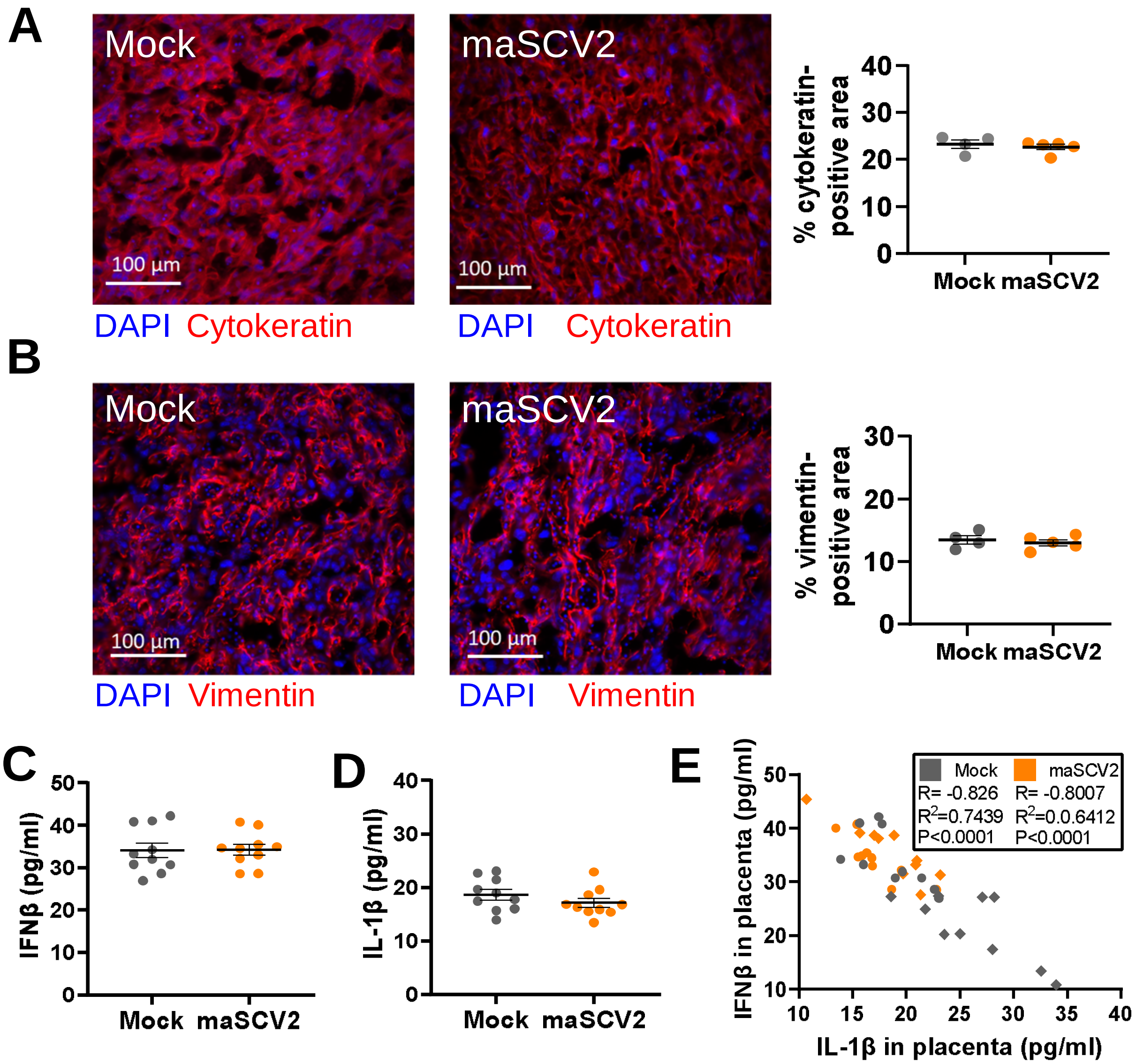


**Supplemental Figure 3. Second trimester-equivalent maSCV2 infection does not result in reduced placental trophoblasts or alter placental IFN-β and IL-1β concentrations.** At embryonic day (E) 10 pregnant dams were intranasally inoculated with $TCID_{50}$ $10^5$ of maSCV2 or mock inoculated with media. At 3 DPI a subset of dams were euthanized to collect maternal and fetal tissues including placentas. Placentas were fixed in 4% PFA and stained or flash frozen. Placentas were immunostained for cytokeratin (A, red) to mark trophoblasts or vimentin (B, red) to mark endothelial cells and DAPI (blue) to label nuclei. Representative images were taken at 20x magnification. Quantification of the percentage positive area for each marker is shown. Flash frozen placentas were homogenized and analyzed by ELISA for IFNβ (C) and IL-1β (D). Associations between IFNβ and IL-1β in placentas of dams infected or mock inoculated at either E10 (circles) or E16 (diamonds) were analyzed by Pearson correlation analyses, with a significant association represented with the R statistic and associated P value (E). Data represent mean +/- standard error of the mean (n= 4-5/group for A-B, each dot indicates 1 placenta and is the mean quantification of 6 fields of view and n=8-10/group for D-E). Significant differences ($p$<0.05) were determined by unpaired two tailed t test.

### Supplemental Figure 3.

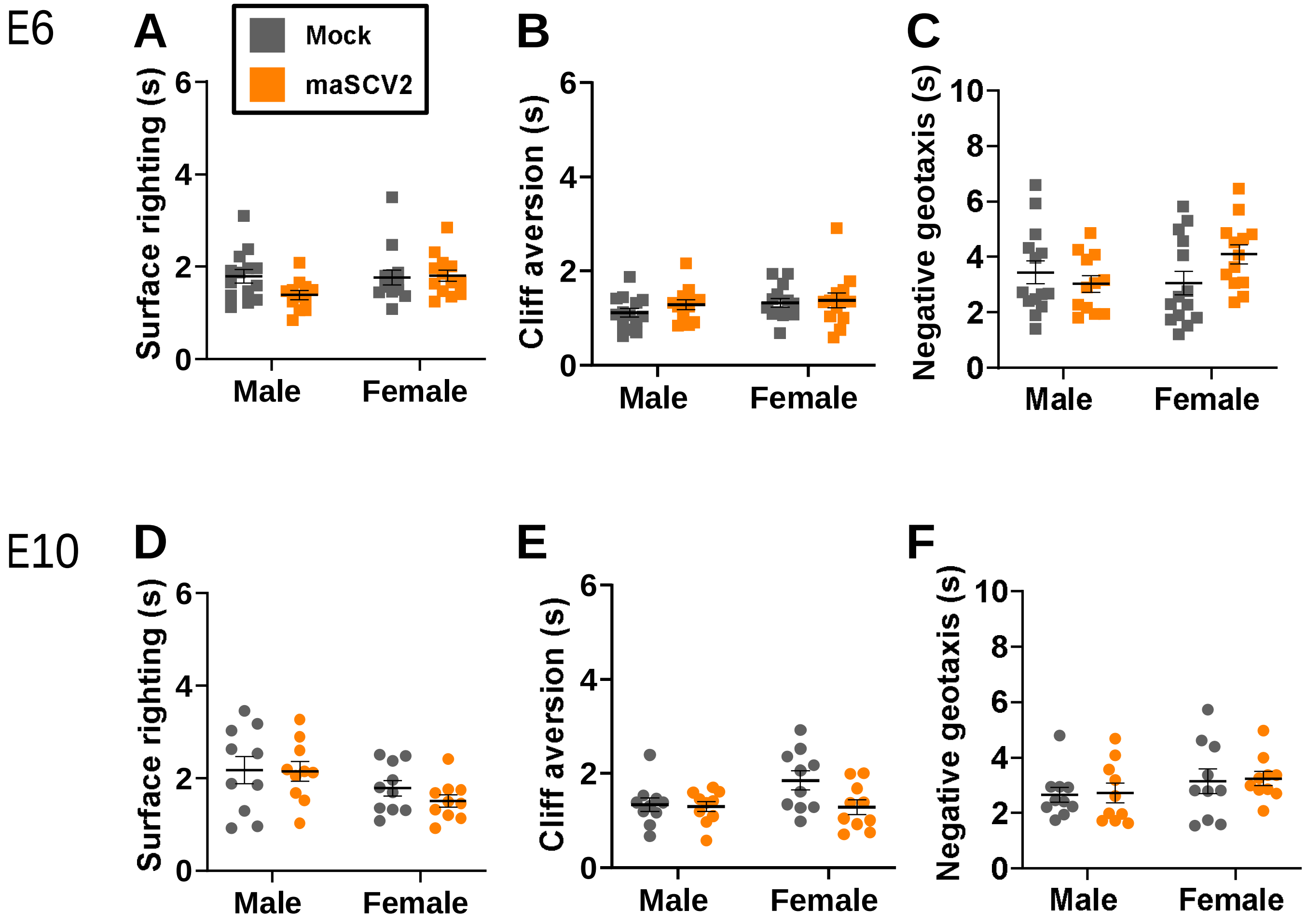


**Supplemental Figure 4. Offspring born to first trimester-equivalent or second trimester-equivalent maSCV2 infected dams do not experience neurodevelopmental delays.**
Pregnant dams were infected at embryonic day (E) 6 or E10 and followed through delivery to characterize adverse offspring outcomes of maternal maSCV2 infection. Offspring were followed to PND5, sexed, and the neurobehavioral assays of surface righting (A,D), cliff aversion (B,E), and negative geotaxis (C,F) were performed to measure neurological development. 1-2 pups per sex per dam were subjected to each test subsequently, with 3 trials given per test, and each pup's best trial for each test was reported (n=7-10 independent litters/group, from two independent replicates). Bars represent the mean ± standard error of the mean with each shape indicating 1 pup. Significant differences ($p < 0.05$) were determined by two-way ANOVA with Bonferroni post hoc test (C-E).

### Supplemental Figure 4.

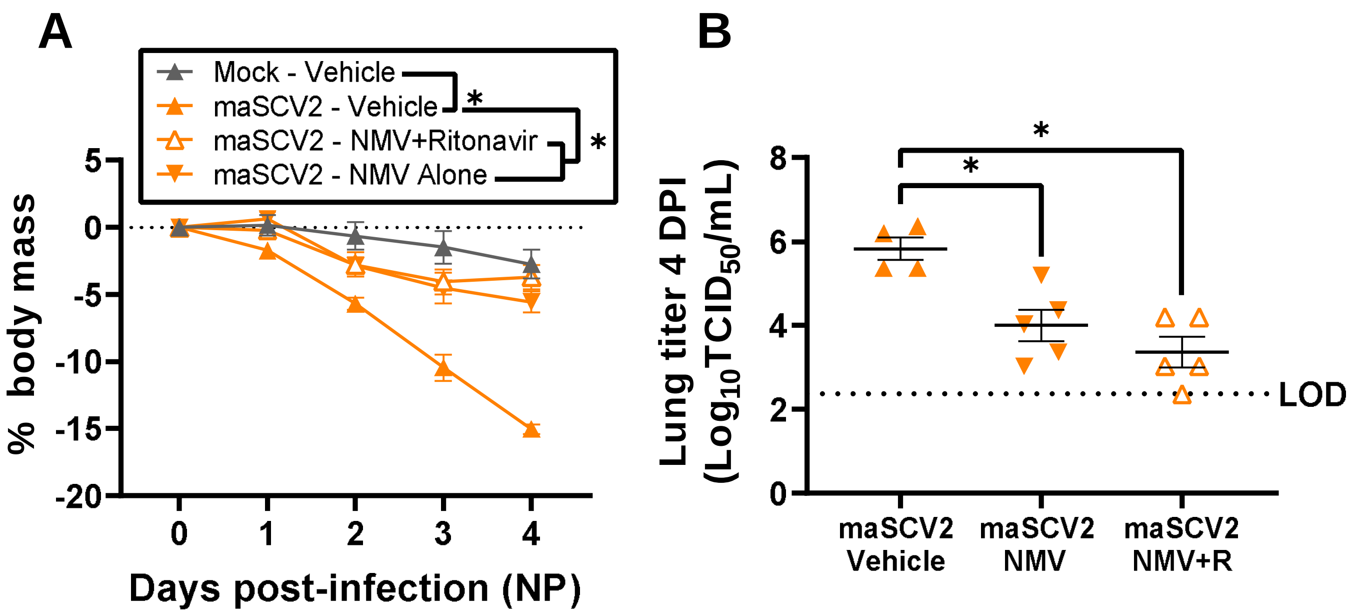


**Supplemental Figure 4. A mouse equivalent dose of ritonavir boosted nirmatrelvir prevents maSCV2 induced morbidity in nonpregnant female mice.** 15-week-old nonpregnant female CD1, which experience greater disease than young adult 8-10 week old nonpregnant female mice, were intranasally infected with $10^5$ $TCID_{50}$ of maSCV2 or mock inoculated with media. Starting at 4 hours post infection and continuing twice daily until 4 dpi were treated with 1.7 mg nirmatrelvir and 0.6 mg ritonavir per dose, 9 mg nirmatrelvir alone (high dose, 300 mg/kg) per dose, or vehicle and were monitored for changes in body mass until 4 dpi (A). At four DPI mice were euthanized for the collection of lung tissue to measure viral titers. Right cranial lungs were homogenized and evaluated for infectious maSCV2 by $TCID_{50}$ assay (B). Bars represent the mean ± standard error of the mean with individual mice indicated by shapes (B; n=4-5/group). Significant differences ($p < 0.05$) were determined by two-way ANOVA with Bonferroni post hoc test of AUCs (A), or one-way ANOVA with Bonferroni post hoc test (B) and are indicated by an asterisk (*). LOD indicates limit of detection.

### Supplemental Figure 5.

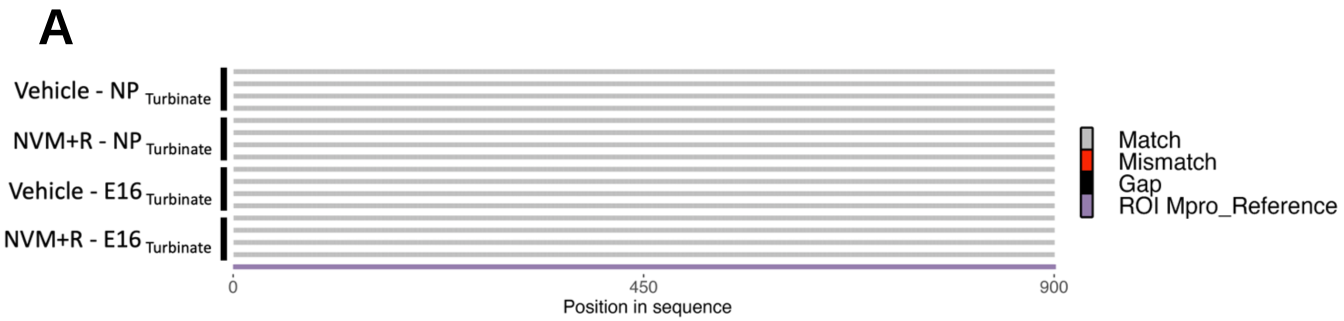


**Supplemental Figure 5. maSCV2 viruses from nasal turbinate tissue of nonpregnant female and pregnant mice infected during the third trimester-equivalent did not acquire mutations associated with antiviral escape.**
At embryonic day (E) 16 pregnant dams or age-matched nonpregnant females were intranasally infected with $10^5$ $TCID_{50}$ of maSCV2 or mock inoculated with media. Starting at 4 hours post infection and continuing twice daily until 3 dpi, mice were treated with 1.7 mg nirmatrelvir and 0.6 mg ritonavir per dose or vehicle. Nasal turbinate tissue was collected at 3 DPI. RNA was extracted from nasal turbinate homogenate, reverse transcribed using ProtoScript® II First Strand cDNA Synthesis Kit, the Mpro region amplified, and Oxford Nanopore sequenced by Plasmidaurus. Consensus sequences were imported and aligned to Mpro using ClustalO v1.2.3 in Geneious Prime v2023.0.4. Alignments were imported into R v4.1.1., visualized, and annotated using seqvisR v0.2.5 (E, n=4/group). Sequence graphic built using Biorender.com.
